# Determining Risk Factors for Triple Whammy Acute Kidney Injury

**DOI:** 10.1101/2021.12.13.472489

**Authors:** Jessica Leete, Carolyn Wang, Francisco J. López-Hernández, Anita T. Layton

## Abstract

Concurrent use of a diuretic, a renin-angiotensin system (RAS) inhibitor, and a non-steroidal anti-inflammatory drug (NSAID) significantly increases the risk of acute kidney injury (AKI). This phenomenon is known as “triple whammy”. Diuretics and RAS inhibitors, such as an angiotensin converting enzyme inhibitor or angiotensin receptor blocker, are often prescribed in tandem for the treatment of hypertension, whereas some NSAIDs, such as ibuprofen, are available over the counter. As such, concurrent treatment with all three drugs is common. The goals of this study are to better understand the mechanisms underlying the development of triple whammy AKI and to identify physiological factors that may increase an individual’s susceptibility. To accomplish these goals, we utilize computational models of long-term blood pressure regulation. These models include variables describing the heart and circulation, kidney function, sodium and water reabsorption in the nephron and the RAS and are parameterized separately for men and women. Hypertension is modeled as overactive renal sympathetic nervous activity. Model simulations suggest that individual variations in water intake, the myogenic response, and drug sensitivity may predispose patients with hypertension to develop triple whammy-induced AKI.

## 1 Introduction

With cardiovascular disease being the leading cause of death in adults worldwide, prescribing safe and effective anti-hypertensive therapies is of great concern. However, a common combination of renin-angiotensin system (RAS) inhibitors, such as angiotensin converting enzyme inhibitors (ACEI) or angiotensin receptor blockers (ARB), with diuretics and easily accessible over the counter non-steroidal anti-inflammatory drugs (NSAIDs) can cause kidney damage. This triple therapy, known as “triple whammy,” was associated with a 31% increased risk for acute kidney injury (AKI), compared to patients treated with diuretic and ACEI/ARB only (95% confidence interval 1.12–1.53) [1]. Triple whammy AKI occurs in 0.88% - 22% of triple treatment patients [1, 2]. AKI may be a serious health condition and has major economic impact. For instance, AKI accounts for 5% of hospital budget [3, 4] and 1% of overall health expenditure [5]. Mortality among AKI patients can be as high as 50-80% of cases in specific populations, such as critically ill patients. Understanding how each of these drugs affect renal autoregulation individually and in combination can help medical providers avoid prescribing dangerous combinations in at risk individuals and administer them with confidence to low risk patients, especially to those who may benefit from pain killers for acute or chronic pain relief.

A goal of this study is to better understand the mechanism by which triple whammy increases the risk of AKI. AKI is marked by higher serum creatinine levels, which indicate a critically low glomerular filtration rate (GFR), according to internationally recognized definitions (i.e. KDIGO) [6] and may be accompanied by low urine flow (less than 400 ml/day) [7]. ACEI and diuretics lower blood pressure through increasing urine flow to reduce blood volume. Combined with NSAIDs, these drugs can prevent the body from properly reacting to hypovolemic states, resulting in dangerously low GFR. In fact, RAS inhibitors and NSAIDs interfere with and thus uncouple the renal afferent and efferent arteriole contractility demand generated by diuretic-induced volume loss to maintain GFR, giving rise to a hemodynamic, pre-renal form of AKI [8]. Triple whammy leads to AKI in some patients but not others. Thus, another goal of this study is to identify which individuals may be particularly susceptible to AKI following triple whammy, and which factors may individually render them so.

To achieve these goals, we apply previously published computational models of blood pressure regulation [9]. Computational models have long been used to describe intereactions between blood pressure regulation and kidney function. In 1972, Guyton and Coleman [10] published their seminal circulation model, concluding that integral to the blood pressure regulation is the pressure-natriueresis curve, whereby higher blood pressure leads to higher sodium excretion in the urine. In 2005, Karaaslan et al. [11] used incorporated renal sympathetic nervouse activity (RSNA) into major components of the Guyton to model to investigate how RSNA contributes to hypertension, and how renal denervation can treat it. In 2014, Hallow et al. [12] extended Karaaslan’s model to include the renin-angiotensin system to investigate efficacy of various antihypertensive therapies. Recognizing that major sex differences exist in blood pressure regulation, Leete and Layton [9] tailored these models to be sex specific. The models were parameterized separately for men and women, and include variables describing circulation, renal function, sodium and water transport through the nephron, and the RAS.

In this study, the mechanisms affected by ACEI, diuretics, and NSAIDs are included and the model is able to directly predict GFR, which is often difficult to obtain experimentally. To identify risk factors for development of AKI, we simulate single, double, and triple drug treatments for normotensive, hypertensive, male, and female humans. Our simulation results reveal a key role of the myogenic response in determining the risk of AKI. Myogenic response, the mechanism by which afferent arteriole resistance is manipulated to maintain appropriate flow through the nephron, is one of the few regulators of GFR untouched by ACEI, diuretics, and NSAIDs. As such, during triple treatment the myogenic response plays a larger than usual role in GFR regulation [8]. We hypothesize that individuals with an impaired myogenic response may be particularly susceptible to triple whammy AKI. Additionally, increased drug sensitivity or low water intake can predispose patients to triple whammy AKI.

## 2 Methods

The present blood pressure regulation models represent the interactions among the cardiovascular system, the renal system, the renal sympathetic nervous system, and the RAS (Fig. 1). A large system of coupled nonlinear algebraic differential equations is used to describe how these systems regulate blood pressure and respond to perturbations. For instance, renal blood flow is adjusted, in part, via renal autoregulatory mechanisms [11, 13] (Eqs. 18 and 19 in the Appendix), according to hormonal and nervous inputs; and renal blood flow, in part, determines Na^+^ excretion, which in turn impacts blood pressure.

**Fig 1.**
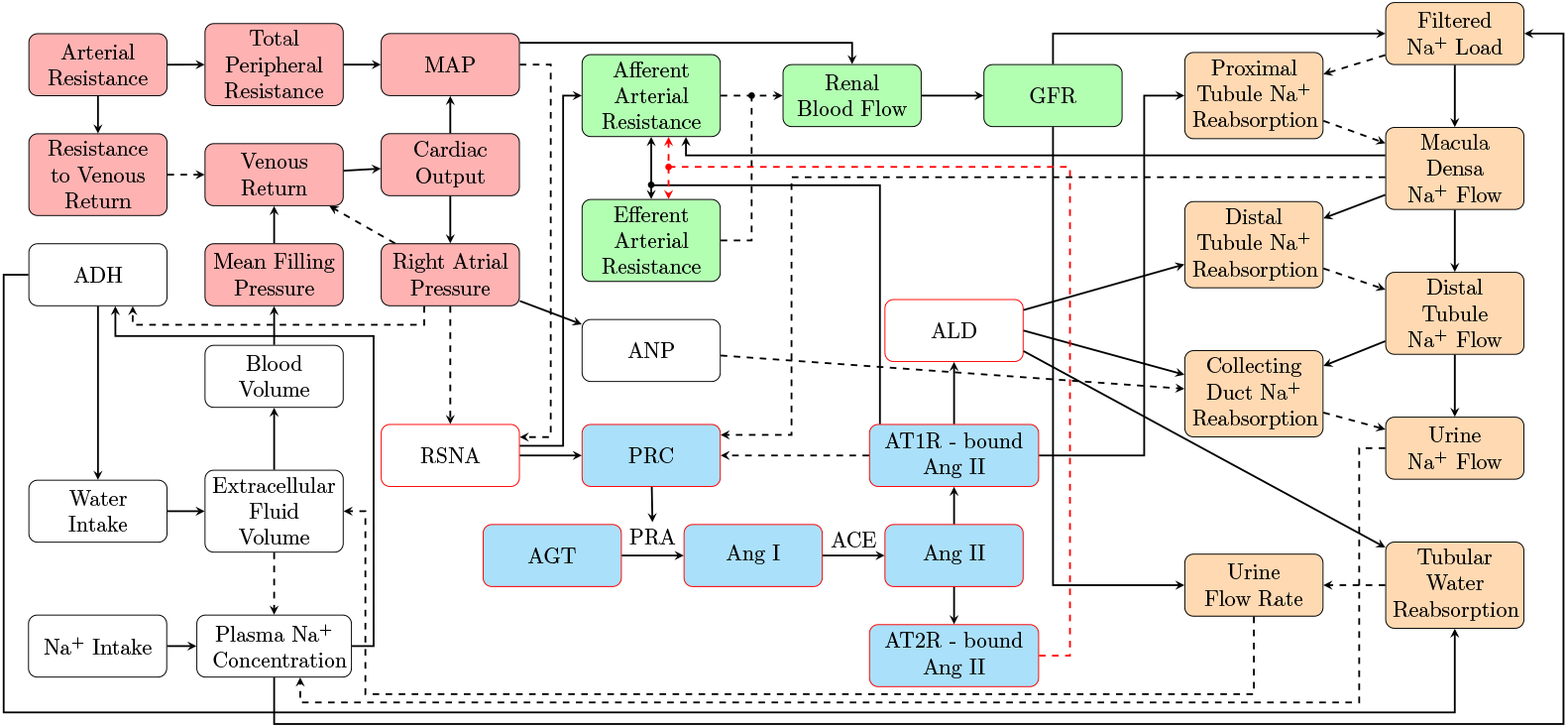
Flowchart of the blood pressure regulation model. Red nodes denote variables describing heart function and blood pressure; green nodes, renal function; orange nodes, Na^+^ and water reabsorption along a nephron; blue nodes, the renin-angiotensin aldosterone system (RAAS). Red outlines describe sex differences between the male and female models.

Below we highlight model components that are particularly relevant to the pharmaceutical treatments considered in this study. Other key model components are described in the Appendix. Unless specified below, model parameters can be found in [9, 11].

### 2.1 Water and sodium intake

An individual’s hypdration status may be a significant determinant in their predisposition to AKI. Water intake (Φ_win_) is dependent on the antidiuretic hormone concentration (*C*_adh_, Eqs. 33-36 in the Appendix)

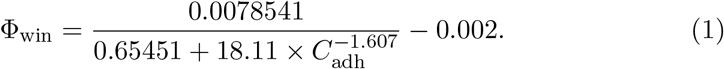

This equation is similar to the one given in [11], but the relationship is modified, in part, to ensure that the model can withstand large increases in urine flow following administration of a diuretic without experiencing extreme hypovolemia.

Low water intake can lead to hypovolemia and low GFR, especially when the mechanisms to enhance renal water retention and to counteract hypovolemia (e.g., via the RAS) are inhibited through drug treatments. As such, in model simulations that consider low water intake, we bound water intake between 0 and 0.001 l/min.

In past models [9, 11, 12], sodium intake is treated as an independent variable, however, for simulations involving changes in sodium excretion, sodium intake is needed to vary to avoid sodium wasting. Sodium appetite is known to vary with aldosterone [14], as such, we allow sodium intake (Φ_sodin_) to gently increase with aldosterone (ALD) concentration (*C*_al_).

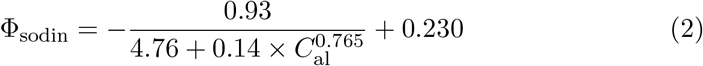

### 2.2 Renal tubular water reabsorption and urine flow

The previous formulations of this model [9, 11, 12] that we are building on are limited in that water and Na^+^ reabsorption rates are independent of each other. This poses a significant problem when one seeks to understand the impact of inhibiting Na^+^ transport (e.g., administration of a diuretic) on urine flow. Other extensions do include a mechanistic formulation of water reabsorption, but require a great level of detail [15]. We choose to balance the need for connection between Na^+^ and water transport without overcomplicating the model for our purpose. Thus, to represent the role of transmembrane Na^+^ concentration gradient in driving water reabsorption, we introduce the term *μ*_Na_ in the tubular water reabsorption equation

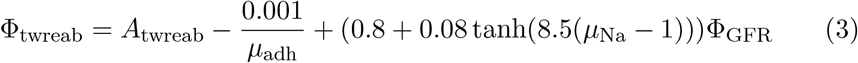

to be

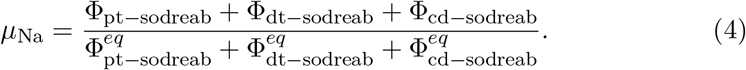

The Φ terms in Eq. 4 represent predicted Na^+^ reabsorption along the three model nephron segments: the “proximal segment (pt)” that includes the proximal tubule and the loop of Henle, the “distal segment (dt)” that includes the distal convoluted tubule and the connecting tubule, and the collecting duct (cd). The superscript “eq” in Eq. 4 denotes the equilibrium Na^+^ reabsorption values. Model parameters are chosen to increase tubular water reabsorption with enhanced Na^+^ reabsorption. *A*_twreab_ in Eq. 3 is the default value of 0.025 as given in [11] unless baseline urine flow falls below 0.001 l/min, then *A*_twreab_ is decreased to yield a baseline value Φ_u_ = 0.001 l/min (specifically, male normotensive, *A*_twreab_ = 0.024791; female normotensive, 0.025; male hypertensive, 0.019178; female hypertensive 0.019959).

Urine flow is given by the difference between GFR (Φ_GFR_) and tubular water reabsorption (Φ_twreab_, Eq. 3).

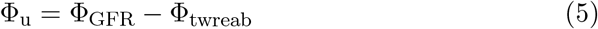

Φ_u_ is bounded above 0.3 ml/min during control simulations to maintain a minimal level of metabolic waste excretion, and we use = 0.27 ml/min or 389 ml/day (consistent with the definition of oliguria [7]) as a threshold to identify AKI.

### 2.3 Blood volume

Blood volume (*V*_b_) is dependent on extracellular fluid volume (*V*_ecf_) and is allowed to decrease so that states of extreme loss of extracellular fluid will cause hypovolemia.

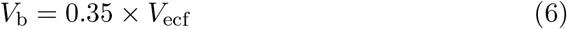

### 2.4 Macula densa feedback

The macula densa cells sense Na^+^ flow entering the distal tubule, and balance SNGFR and thick ascending limb transport capacity via the tubuloglomerular feedback (TGF). When the macula densa cells sense a below-target Na^+^ flow, TGF is activated to dilate the afferent arteriole, thereby increasing renal blood flow and SNGFR; and vice versa (Fig. 2A). The model afferent arteriole resistance (*R*_aa_) is proportional to Σ_TGF_ (Eq. 18), which denotes the feedback effect of macula densa Na^+^ flow (Φ_md−sod_, Eq. 17). TGF is affected by both furosemide and NSAID treatment, represented by the term *κ*_f−md_ and the indicator function, 𝕀 (), respectively, in Eq. 2. How the effects of these two drugs are fitted is discussed in later sections.

**Fig 2.**
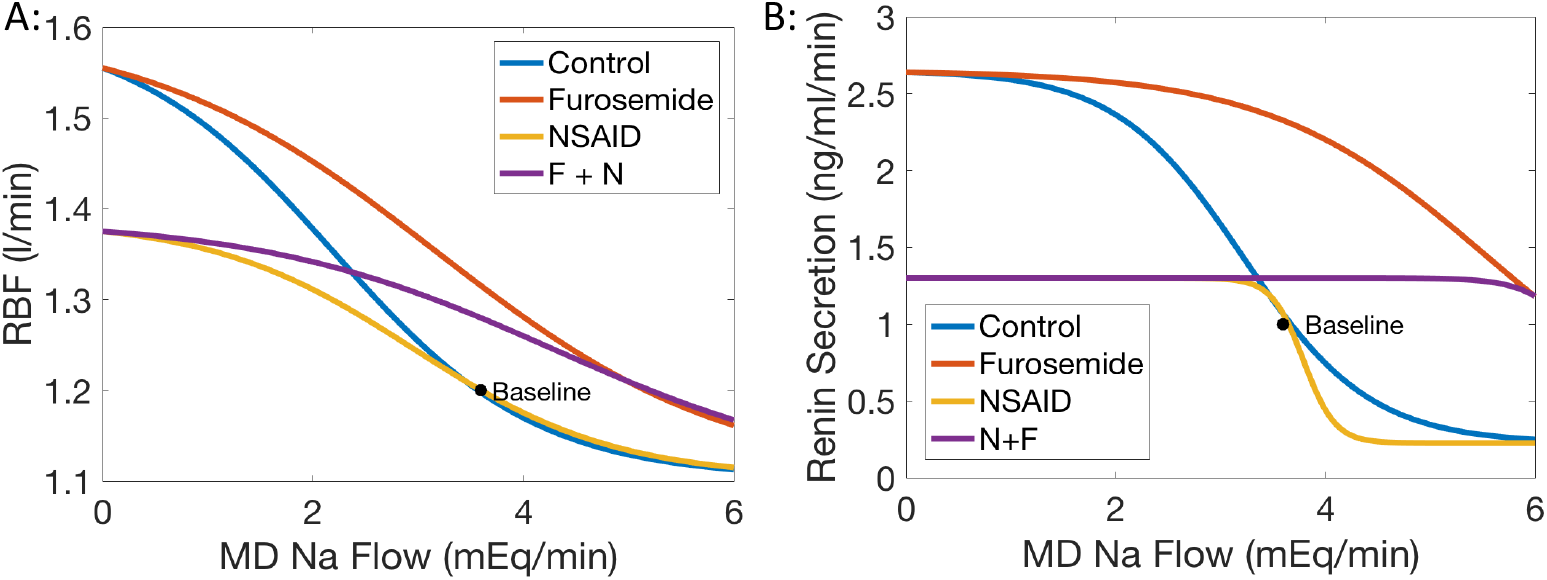
Effects of furosemide and NSAIDs on renal blood flow (RBF) and renin secretion as functions of macula densa Na^+^ flow. For each panel, macula densa N^+^ flow is treated as the independent variable and the changes in (A) RBF and (B) renin secretion are tracked assuming all other factors are held constant. (A) RBF decreases at higher macula densa (MD) Na^+^ flow via the tubuloglomerular feedback (TGF), obtained for four treatment conditions: control (no drug), furosemide only, NSAIDs only, and furosemide and NSAIDs (F + N). The effects of furosemide and NSAIDs are represented in the TGF function Σ_TGF_ (Eq. 7). (B) Renin secretion decreases at higher MD Na^+^ flow. That relation, and how it is affected by the administration of furosemide and NSAIDs, is represented in the feedback function *ν*_md−sod_ (Eq. 8).

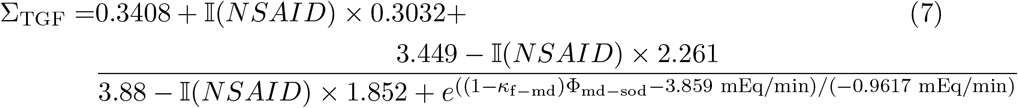

Another feedback mechanism is one in which low macula densa Na^+^ flow triggers increased renin secretion to activate the RAS and to enhance Na^+^ retention, and vice versa (Fig. 2B). Renin secretion (*R*_sec_) is proportional to *ν*_md−sod_, which is the feedback effect of macula densa Na^+^ flow (Φ_md−sod_, Eq.17). This feedback effect is changed by both furosemide and NSAID treatment, represented by the term *κ*_f−md_ and the indicator function, 𝕀(), respectively:

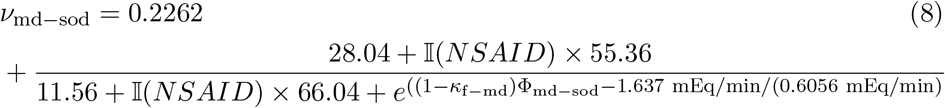

Descriptions of the effect of NSAIDs and furosemide on equation 8 are found in sections 2.7 and 2.8.

### 2.5 Simulating hypertension

Hypertension is a multifaceted disease. The scope of the present study is limited to hypertension that is induced by an over-active RSNA as done in [9]. That is represented by increasing the baseline RSNA parameter (*N*_rsna_, see Eq. 12 in the Appendix) by 2.5-fold. This results in constriction of the afferent and efferent arterioles, stimulation of the RAS, and enhanced renal Na^+^ reabsorption.

### 2.6 Simulating administration of ACEI

To simulate the effect of ACEI, we reduce the rate at which Ang I is converted to Ang II, denoted *c*_ACE_ (see Eq. 27), by a target level *κ*_ACEI_:

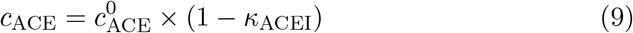

where 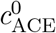 is the baseline Ang I-to-Ang II conversion rate, and 0 ≤ *κ*_ACEI_ < 1. To determine the value of *κ*_ACEI_, we compared our model to experimental data. Delles et al. [16] found that after a week of treatment in hypertensive male patients, ACEI treatment with 20mg/day of enalapril reduced supine mean arterial pressure (MAP) by 11% and did not change GFR. To replicate these changes in MAP and GFR in the steady state models, we use *κ*_ACEI_ = 0.76 for all models.

### 2.7 Simulating administration of furosemide

Furosemide, a loop diuretic, inhibits the Na^+^-K^+^-Cl^−^ cotransporter 2 (NKCC2) along the thick ascending limb of the loop of Henle. In our model, this is included in the Na^+^ reabsorption of the model proximal segment, which consists of the proximal convoluted tubule, S3 segment, and the loop of Henle. To model the inhibition of NKCC2, we reduce fractional model proximal segment Na^+^ reabsorption (*η*_pt−dosreab_) by a percentage *κ*_f_, 0 ≤ *κ*_f_ < 1. As such, Eq. 14 becomes

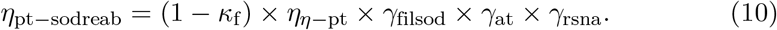

Furosemide also inhibits the NKCC2 transporters in the macula densa, which interferes with macula densa Na^+^ signaling and has implications on TGF signal and renin secretion. We describe the inhibition of Na^+^ sensing in the macula densa cells by *κ*_f−md_, where 0 ≤ *κ*_f−md_ < 1. By multiplying Φ_md−sod_ by (1 − *κ*_f−md_) in Eqs. 7 and 8, we model the effect of NKCC2 inhibition by furosemide such that the macula densa cells only sense a fraction of the Na^+^ flowing past in the lumen.

Experimental treatment with 40 mg i.v. furosemide results in a 70% increase in ALD [17], while a 0.5mg/kg mg bolus dose followed by 0.5 mg/kg per hour i.v. caused a 12% decrease in GFR and a 10-fold increase in urine flow around an hour after starting treatment [18]; these studies did not separate men and women. In another study, plasma renin activity (PRA) for men and women were reported to increase by 132% and 117% increase, respectively, when given a 40 mg dose intravenously [19]. While these exact changes could not be matched, *κ*_f_ = 0.08, *κ*_f−md_ = 0.3 were chosen to be used in all models to increase urine flow, ALD, and PRA, as well as decrease GFR relative to steady-state control.

### 2.8 Simulating administration of NSAIDs

NSAIDs inhibit the COX-1 and COX-2 enzymes, which play a key role in macula densa Na^+^ signaling pathway during low Na^+^ flow. The mechanisms that are affected by NSAIDs include TGF and a feedback effect on renin secretion.

According to Araujo et al. [20], COX-2 inhibition reduces TGF by 46%. To match this, in Eq. 7 with NSAID treatment and during low Na^+^ flow, the change in RBF is reduced by 46% (Fig. 2A).

In a study done by Araujo et al. [20], COX-2 inhibition reduced the maximum change in proximal tubule stop-flow pressure caused by TGF during different proximal tubule perfusion rates by 46%. To match this, in Eq. 7 with NSAID treatment, the maximum change in afferent arteriole resistance during low macula densa sodium flow is reduced by 46%. This causes a reduction in the increase in RBF when all other factors are held constant (Fig. 2A).

Traynor et al. [21] found that with COX-2 inhibition, the increased renin secretion in response to low lumen sodium flow is eliminated. Thus, with NSAID treatment, Eq. 8 is designed to level off during low Na^+^ flow, stopping renin secretion from increasing (Fig. 2B).

### 2.9 Simulating impaired autoregulation

TGF and myogenic activity accounts for 90% of renal autoregulation [22]. Under double treatment with ACEI and furosemide, the body’s ability to defend against hypovolemia through the RAS and renal sympathetic nervous activity (RSNA) is eliminated and regulation of GFR falls to TGF and the myogenic response [8]. Hence, we assess the extent to which patients who have impaired myogenic regulation are at increased risk for double or triple whammy AKI. To investigate this, we describe an impaired myogenic effect using the following formula to link glomerular hydrostatic pressure to afferent arteriole resistance (Eq. 18):

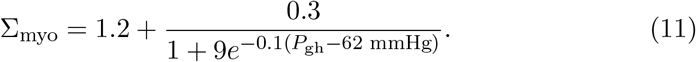

Compared to the analogous formula for normal myogenic effect (Eq. 21), the above formula attenuates the reduction (rise) in afferent arteriole resistance during low (high) levels of glomerular hydrostatic pressure, thereby narrowing the range of blood pressure within which GFR is stable.

## 3 Results

The model is run forward in time for 2 simulated days. To simulate the administration of ACEI or furosemide, the relevant parameters (*κ*_ACEI_, *κ*_f_, *κ*_f−md_) are gradually increased over the initial 30 minutes from the baseline values to their target values. Average GFR, MAP, and PRA as well as total urine volume for each day are then calculated.

### 3.1 Response to hypertensive stimuli

Key model predictions [urine volume, GFR, MAP, PRA] are shown in Fig. 3. The simulations without drug treatment are labelled “Control.” Note that in control simulations, the MAP increase predicted for the hypertensive female model is lower than the hypertensive male model. Our model includes sex differences in RSNA, in that the female model has less excitable and more easily repressed RSNA [9, 23]. This leads to a smaller actual increase in female RSNA, and thus a smaller increase in MAP (Fig, 3a).

**Fig 3.**
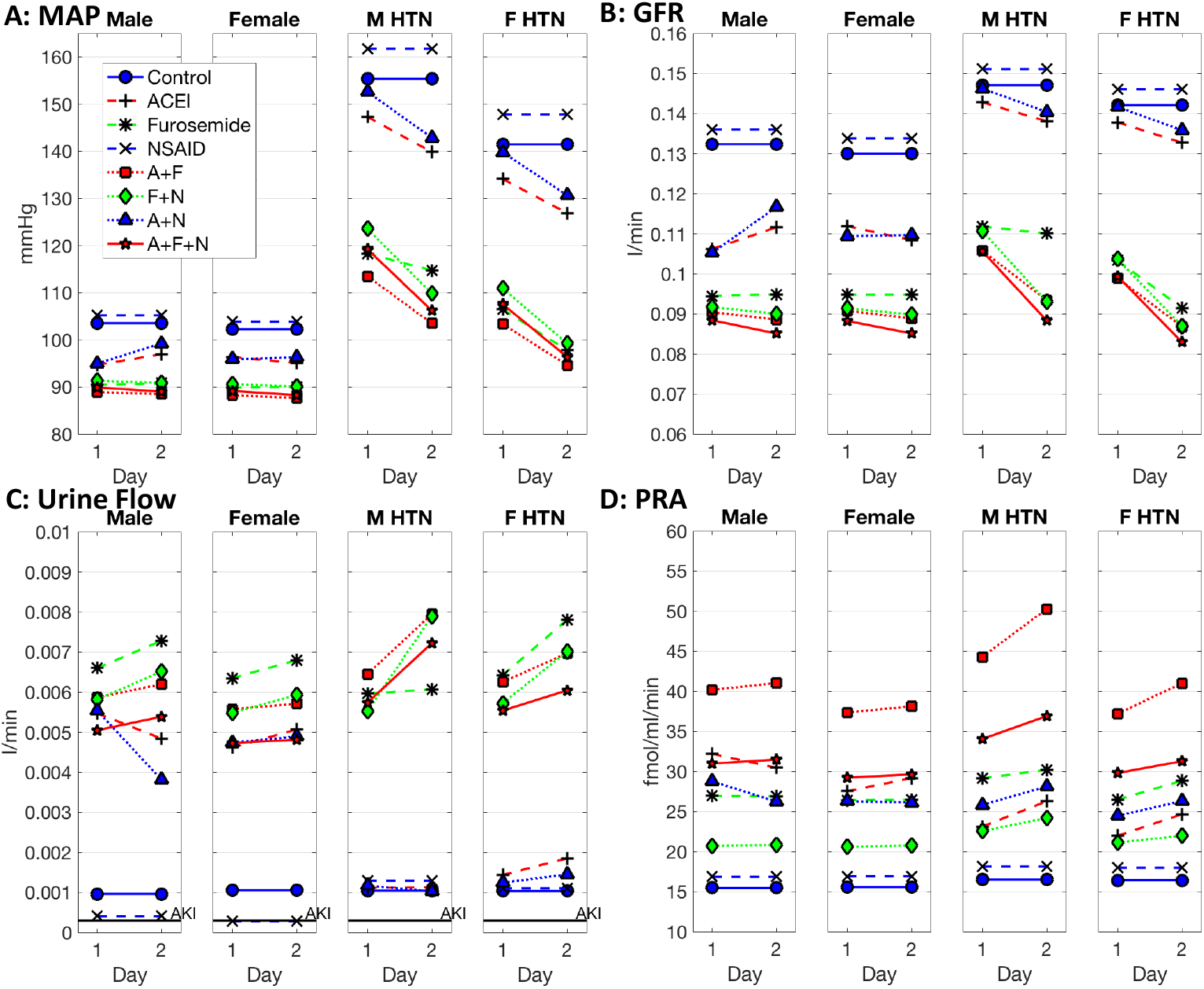
Effect of drug treatments on renal and cardiovascular function. Shown are key model predictions for male normotensive (“Male”), female normotensive (“Female”), male hypertensive (“M HTN”), and female hypertensive (“F HTN”) models, obtained under control conditions (i.e., no drug) and following single, double, and triple drug treatments. (A) mean arterial pressure (MAP); (B) glomerular filtration rate (GFR); (C) urine volume; (D), plasma renin activity (PRA). Two data points are computed for each case, corresponding to key variables averaged over days 1 and 2. Triple treatment yields the lowest GFR (B), albeit not to a level indicative of acute kidney injury.

In our model, a higher MAP raises GFR in control simulations (Fig. 3). Specifically, the 44.3 and 33.2 mmHg increases in MAP predicted in the male and female hypertensive models are associated, respectively, with 12.4% and 9.4% increases in GFR. In contrast in population studies, higher daytime diastolic blood pressure is correlated with lower GFR [24]. This paradoxical effect may be attributable to the renal damage caused by elevated blood pressure, which is not captured in our hypertensive models.

### 3.2 Response to drug treatments

Key model predictions for single drug treatments are shown in Fig. 3, labelled “ACEI”, “Furosemide,” and “NSAIDs.” ACEI and, to a larger extent, furosemide lower MAP and GFR, and those decreases are larger in hypertensive models than in normotensive models. For both normotensive and hypertensive models, all single drug treatment simulations yield urine volume substantially above threshold for AKI. Daily averages were quantitatively similar to end of day measurements, with a few exceptions. While clinical measurements often only capture data at a single point in time, looking at daily averages offer a more detailed look at model behavior.

Parameters describing ACEI treatment were fitted to match the change in GFR and MAP in hypertensive patients during chronic treatment in Delles et al. [16]. PRA has been observed to increase by 90% at the end of three weeks of 5mg daily ramipril in mild hypertensive patients [25]. Male and female hypertensive model simulations show a 66.6% and 52.5% percent increase in PRA, respectively, on average during the second day (Figure 3). Model simulations show a slight increase in the plasma Na^+^ concentration (146.5 mEq/L versus 144 mEq/L at baseline) in both the male and female normotensive models, indeed, hyponatremia is a very rare complication of ACEI [26].

Parameters describing furosemide treatment were fitted to match trends in urine flow, ALD, PRA, and GFR in chronic treatment [17–19]. Model simulations show a 8.9 mmHg and 9.9 mmHg drop in MAP in normotensive male and females, consistent with the 10 mmHg drop found by [27] during during acute treatment of 40 mg in normotensive men. The drop in MAP is driven by increases in urine volume. Normotensive model results predict a 24 hour urine volume of 4.2 L and 3.7 L (194.3% and 110% increase) in male and female respectively during the second day. These values are within the range found by Hutcheon et al. [28] who observed a 122% and 205% increase in 12 hr urine volume for 100 mg and 200 mg single oral dose in healthy individuals but is lower than Hammarlund et al. [29] who found 2-2.2 L per 8 hrs for 40 mg oral or i.v. dose. Hutcheon et al. [28] also found no change in plasma Na^+^ concentration; model simulations show a small increase of 1.3 and 0.77 mEq/L for male and female normotensive, respectively. Gottleib et al. [30] found that chronic heart failure patients saw a 25% decrease in GFR in the first two hours after furosemide administration. Our models show a 22.3% and 22% decrease in GFR in normotensive male and females, respectively (26% and 28% for hypertensive models). Our normotensive models show an increase in PRA (50.8% male and 46% female) while Haber et al. [31] found an 43.75% increase from 80 mg furosemide compared to control with low Na^+^ intake, and a much lager increase (386%) compared to control with high Na^+^ intake in healthy individuals. Kelly et al. [27] found a 130% increase in Ang II while our models predict a 50.7% and 46% in male and female normotensive models respectively. They also reported a 71% in plasma ALD while our models predict a 121.6% and 126.4% for male and female normotensive respectively.

Experimentally, NSAID treatment causes little or no increase in MAP in normotensive patients, whereas an increase in MAP has been reported in hypertensive patients (3.32 mmHg [32]; 5mmHg [33, 34]). However, that increase in MAP does not come through water and Na^+^ retention [34] but rather through an increase in arterial pressure and peripheral resistance and possible decline in cardiac output [35]. NSAIDs have also been observed to lead to reductions in PRA and ALD [33]. The present models do not predict the drop in PRA (Fig. 3D) or ALD (results not shown), nor do they predict a increase in MAP following NSAID treatment alone (Fig. 3A).

We then consider combination treatments involving pairs of drugs. Key results are shown in Fig. 3. ACEI and furosemide together create a stronger MAP lowering effect in all models compared to either single treatment. In contrast, the addition of NSAIDs does not have a significant effect on MAP lowering effect of furosemide in any model. None of the double treatments reduces urine volume below the AKI threshold.

Triple treatment involving ACEI, furosemide, and NSAIDs results in MAP levels similar to double treatment with ACEI and furosemide. Hypertensive simulations are above the urine threshold for AKI, but normotensive simulations show low urine volume on the second day (Fig. 3C). A slightly larger decrease in GFR is predicted for the hypertensive models (31.7% and 33.9% in male and female, respectively) than the normotensive models (30.6% and 30.2% in male and female, respectively), stemming from a larger drop in MAP in the hypertensive models. However, GFR drops to slightly lower levels in the female hypertensive model. Taken together, none of the models predict critically low GFR following triple treatment. This result may not be unexpected, given that only a fraction (0.8%-22%) of patients subjected to triple treatment actually suffer pre-renal azoteamia stemming from low GFR [1, 2]. As such, a hypertensive patient without additional autoregulatory problems is unlikely to suffer from AKI following triple treatment.

### 3.3 Potential risk factors for AKI

We conduct simulations to assess the effects of potential risk factors on the predicted response to double and triple treatments. The risk factors considered are impaired renal autoregulation, low water intake, and high drug sensitivity. Impaired renal autoregulation is simulated by attenuating the myogenic response (Eq. 11). Low water intake is simulated by bounding water intake to below control level (0.001 l/min). Increased drug sensitivity is simulated by increasing the parameters associated with ACEI and furosemide: *κ*_ACEI_ = 0.9, *κ*_f_ = 0.3, *κ*_f−md_ = 0.5 (compared to standard values of 0.76, 0.08, and 0.3, respectively), and we further assume that NSAIDs completely inhibit TGF. Predicted MAP and GFR at the end of the second day as well as urine volume during the two days are shown in Fig. 4 for the male and female hypertensive models.

**Fig 4.**
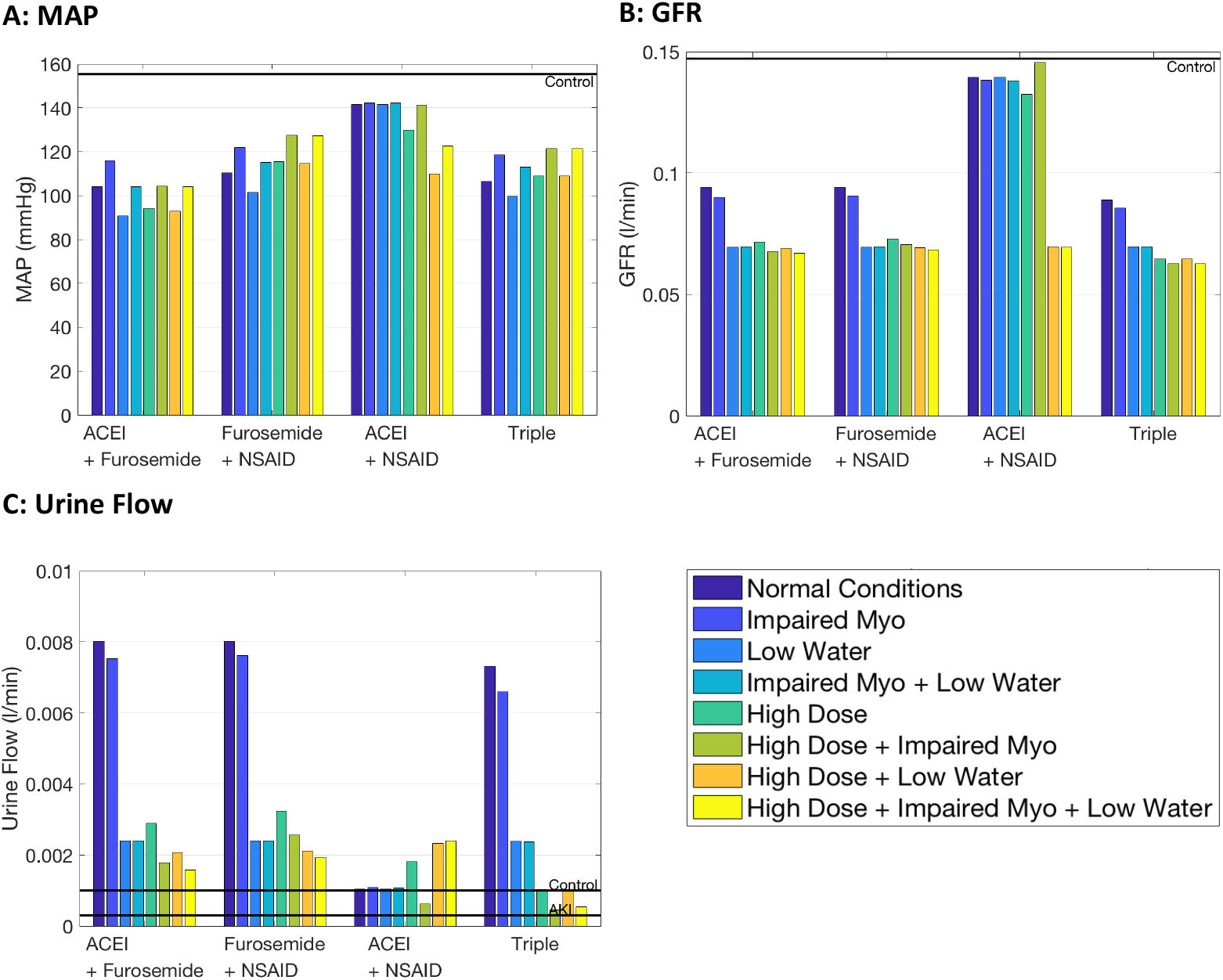
Effect of selected drug combinations on renal and cardiovascular function of the male and female hypertensive models, obtained under differing aggravating conditions. (A) mean arterial pressure (MAP) and (B) glomerular filtration rate (GFR) values at the end of the second day of treatment as well as (C) urine volume over two days of treatment. Triple treatment yields the lowest GFR, albeit not to a level indicative of acute kidney injury. Solid lines labelled “Control” indicates values without drug treatment. Triple treatment administered to a highly sensitive patient yields the lowest urine volume (C).

Recall that in the absence of these risk factors, ACEI and furosemide together lower GFR (by 34.5% and 38.1% in hypertensive male and female models, respectively) further than single treatments or double treatments with NSAIDs (Fig. 3B). When the effect of ACEI and furosemide is not counteracted by a robust myogenic response, GFR is further reduced (by 37.6% and 39.6%). When further aggravated by low water intake, GFR drops even lower (by 51.3% and 47.9%). It is noteworthy that in the presence of a robust myogenic mechanism, low water intake significantly lowers GFR.

In any double or triple treatment with ACEI or furosemide, low water intake reduces urine volume far more than can be accounted for by reduced GFR alone (Fig. 4). During treatment under normal conditions, GFR decreases because MAP decreases. Low water intake exacerbates this effect. During low water intake, tubular water reabsorption still decreases overall, but less than the decrease in GFR due to an increase in ADH (results not shown). As such, when comparing low water intake to normal conditions, GFR decreases further than water reabsorption, causing a more drastic decrease in urine flow.

An impaired myogenic response has a minimal effect on GFR either alone or in addition to another potential risk factor. However, during triple treatment with high sensitivity to drugs and low water intake, myogenic response impairment does decrease urine volume. In this situation, the decrease in GFR is not met with a matching decrease in tubular water reabsorption; in fact, water reabsorption does not change with low water intake during high sensitivity to triple treatment (results not shown). This is in contrast to most other simulations where a decrease in GFR is accompanied by reduced tubular water reabsorption due to the direct dependence on GFR (Eq. 3).

Somewhat surprisingly, high drug sensitivity does not lead to further drops in MAP in the case of furosemide + NSAID, ACEI + NSAID, or triple treatment. However, a higher dose of ACEI + furosemide does increase the MAP lowering effect. Triple treatment with high drug sensitivity is predicted to lower GFR to 0.061 and 0.056 l/min in male and female hypertensive models, respectively, which correspond to 29.9% and 29.3% reduction from normal conditions (i.e., triple treatment at regular sensitivity). Triple treatment with high drug sensitivity is also where the model experiences the lowest urine volume, especially in the female hypertensive model. Small decreases in GFR can translate to large decreases in urine volume, especially when the decrease in GFR is not matched by a similar decreases in tubular water reabsorption, as in this case. Excessive sensitivity to drugs may lead to increased risk for AKI; this finding is especially important as many dosages are only based on clinical trials including men, rather than finding appropriate dosages for women separately.

The male and female hypertensive models act similarly in most situations, the exception being ACEI and NSAID double treatment. Here the female hypertensive model shows that GFR is either maintained or drops to a similar low level. GFR regulation has a plateau, where GFR is maintained at a constant value for a range of MAP. The plateau is narrower now due to drug treatments or potential risk factors breaking these feedback systems. Combined with the lower MAP due to treatment or low water intake, the system is pushed out of the regulatory plateau range. Small variations between the male and female models may change exactly where the edge of the regulatory plateau resides.

## 4 Discussion

AKI is typically associated with elevated serum creatinine levels and a severe drop in urine output. The present study is concerned with “pre-renal AKI,” which results from altered renal hemodynamics leading to critically lowered GFR. (“Renal AKI” is caused by drug or ischemia-induced parenchymal injury, whereas “post-AKI” is a result of urinary tract obstruction.) Pre-renal AKI may develop as a result of hypotension, dehydration, heart failure, renal microangiopathy, or following the administration of certain drug treatments. Triple treatment with ACEI, diuretics, and NSAIDs is known to increase the rate of AKI [1]. However, the factors that predispose some patients to the development of triple whammy AKI are incompletely characterized. Our study has identified dehydration and high sensitivity to drug treatment as key contributing factors to the development of triple whammy AKI.

To maintain normal kidney function, GFR is kept within a narrow range and exhibits a wide plateau for renal perfusion pressures 80–180 mmHg [36]. A key autoregulatory mechanism is the myogenic response, which is a property of the preglomerular vasculature wherein a rise in intravascular pressure elicits constriction that generates a compensatory increase in vascular resistance.Another autoregulatory mechanism is TGF, which is a negative feedback response that balances glomerular filtration with tubular reabsorptive capacity. Both mechanisms operate via regulation of the contractility of the afferent arteriole smooth muscle cells.

In TGF, the macula densa cells located at the end of the cortical thick ascending limb act as the salt sensor. Tubular salt sensing by the macula densa involves apical NaCl transport mechanisms, including the furosemide-sensitive Na^+^-K^+^-2Cl^−^ cotransporter (NKCC2), which is the primary NaCl entry mechanism. When tubular fluid [Na^+^] and [Cl^−^] at the macula densa exceed their respective target values, the macula densa cells signals to constrict the afferent arteriole, and vice versa. Among the mediators produced by the macula densa cells are the vasodilator prostaglandins, which are derived from the COX-2 pathway and are critical in the TGF mechanism and renin signaling cascade. Renin activates the systemic and renal RAS, which increase blood pressure and GFR.

RSNA regulates blood pressure via the natriuresis-diuresis-driven volemic response, and GFR via afferent and efferent arteriolar tone as well as proximal Na^+^ reabsorption. Specifically, the RSNA-mediated responses include the (i) release of ADH, which increases GFR by inducing afferent vasodilation and efferent vasoconstriction, (ii) renin secretion, which activates RAS and raises blood pressure, and (iii) enhanced Na^+^ reabsorption, leading to volume expansion and blood pressure increases [37–39].

Yet another factor in the interplay between blood pressure and GFR is blood volume. Perturbations in blood volume stimulate the stretch/distension receptors located on the right auricle and central veins, and induce responses aimed at stabilizing blood pressure. An increase in blood volume gives rise to atrial stretch and activates the release of atrial natriuretic peptide from cardiomyocytes. Atrial natriuretic peptide induces afferent vasodilation and efferent vasoconstriction; consequently, GFR increases, followed by natriuresis and diuresis.

### 4.1 Response to drug treatments

The individual effect on GFR and blood pressure of each of the above mechanisms is well characterized. However, how they interact, especially under drug-induced alterations, is not completely understood in an integrated sense. Diuretics, for instance, lowers extracellular fluid volume by inhibiting tubular solute and water reabsorption and inducing natriuresis and diuresis. Taken in isolation, the reduction in extracellular fluid volume leads to reduction in blood volume and blood pressure, especially in hypertensive patients (Fig. 3A). ACEI inhibit the synthesis of Ang II, a potent vasoconstrictor and anti-natriuretic peptide that promotes blood pressure elevation. Another commonly prescribed RAS inhibitor is the ARB, which impedes the binding of Ang II to Ang II type I receptor (AT1R), which is implicated in vasoconstriction, ALD secretion, and enhanced tubular salt and water reabsorption. ACEI and ARB lower blood pressure, especially in hypertensive patients (Fig. 3A), but alone do not significantly alter GFR (Fig. 3B). NSAIDs inhibit the production of prostaglandins, which, under hypovolemic conditions, may have an adverse effect on salt and water retention, and GFR regulation.

As previously noted, a number of blood pressure compensatory mechanisms are activated following the administration of diuretics (e.g., RSNA and RAS). The effects of these mechanisms are significantly attenuated by the concurrent administration of an ACEI (or ARB). As a result, model simulations predict that the combined administration of ACEI and furosemide slightly lower blood pressure and GFR, compared to furosemide alone, in normotensive populations; the predicted blood pressure and GFR reductions are more significant in hypertensive populations (Figs. 3A and 3B). The competing effects of lower GFR and lower tubular Na^+^ and water reabsorption results in the attenuation by ACEI of the diuretic effect (i.e., urine output) of furosemide in normotensive models and the female hypertensive model but increase the diuretic effect of furosemide in male hypertensive model (Fig. 3C). Some experimental findings have revealed that ACEI attenuate the diuretic effect of furosemide in normotensive humans [17] and normotensive Sprague-Dawley rats [40] while others have found that in chronic heart failure patients ACEI and furosemide double treatment increased urine flow in comparison to furosemide only treatment [41]. Thunhorst and Johnson [40] found the dosage of ACEI to be the determining factor: low dosage of captopril was found to increase urine flow, whereas a high dose decreased urine flow, possibly due to the RAS being a mediator of water ingestion. This may suggest that dosing needs to be tailored to the individual in order to achieve the intended diuretic effect and properly reduce blood pressure or to avoid problematic side effects.

NSAIDs have been consistently found to attenuate the diuretic and RAS activating effect of furosemide in normotensive and hypertensive humans [33, 40, 42] and normotensive Sprague-Dawley rats [43]. Simulation results are in general agreement with these findings. In the models, urine volume is predicted to be attenuated with NSAID and furosemide double treatment compared to furosemide single treatment, with the exception of the male hypertensive model where it increases with double treatment (Fig. 3C). The increase in PRA from furosemide is also attenuated with NSAID treatment (Fig. 3d). These attenuations of renin and urine volume stems from NSAIDs inhibition of macula densa sodium signaling. Diuretic attenuation occurs due to the decrease of GFR from TGF (Fig. 2A), while RAS attenuation occurs to the inhibition of macula densa signaling of renin secretion (Fig. 2B). As such, this double treatment may put patients at risk for AKI as the RAS’ ability to respond to a hypovolemic state may be impaired.

ACEI and NSAIDs together have been shown to not decrease blood pressure and GFR in normotensive individuals [44] and to have no effect on RBF and GFR in ACEI treated hypertensive patients with controlled blood pressure [45]. However, the renin increase due to ACEI is attenuated with COX-2 inhibition [46]. While our hypertensive models predict a decrease in MAP in response to ACEI single treatment, the addition of NSAIDs slightly weakens that response (Fig. 3A). Model predictions differ from experimental results in that the renin increase from ACEI is somewhat enhanced with NSAIDs (by ∼18%) in the hypertensive models (Fig. 3D). In a retrospective cohort study, Lapi et al. [1], found that double treatment with ACEI and NSAIDs is not associated with an increased risk of AKI. Consistent with that observation, model simulations predict similar urine volume levels in all models (Fig. 3C). Thus, no indication of an elevated risk for AKI is predicted following the combination therapy of ACEI and NSAIDs, in the absence of additional risk factors.

In triple treatment with ACEI, diuretics, and NSAIDs, critical blood pressure and GFR regulatory mechanisms are simultaneously interrupted. As previously noted, diuretics decrease extracellular fluid volume and blood pressure. The renal autoregulatory mechanisms that maintain GFR via TGF- and RAS-mediated compensation are inhibited through ACEI and NSAID. Consequently, triple therapy has been reported to increase the rate of AKI [1, 47–49]. Perhaps not unexpectedly, model simulations indicate that triple treatment reduces GFR more than single or double treatments in all individuals (Fig. 3B). However, under triple treatment, urine volume and GFR have not been predicted to fall sufficiently far to indicate AKI. This result is consistent with the fact that only a fraction of individuals develop AKI following triple treatment. Thus, we expect individuals who may be hypertensive but are otherwise healthy to be able to withstand triple treatment—in the absence of other aggravating factors.

### 4.2 Potential risk factors for AKI

One aggravating factor explored in the present study is high drug sensitivity, which, perhaps not unexpectedly, can substantially lower GFR and urine flow (Fig. 4) through further inhibiting the feedback mechanisms discussed previously. Differences in pharmacokinetics and metabolism can affect the drug concentrations at target sites, the effect of each drug, and thus increase the risk for AKI in some individuals.

We also considered the effect of an impaired myogenic response, as seen in cardiovascular and renal diseases [50, 51]. As previously noted, GFR normally exhibits a wide plateau for a range of renal perfusion pressure values. That plateau can be attributed, in large part, to the myogenic response; by comparison the contribution of TGF is significant only for a narrow range of pressure values around the baseline [52]. When renal autoregulation is impaired, GFR becomes more sensitive to variations in renal perfusion pressure, which drops following the administration of diuretics and ACEI. An excessive reduction in GFR may result in AKI. Normally, GFR needs to decrease beyond 60% for plasma creatinine to significantly increase, and thus for AKI to ensue, due to the increased creatinine secretion that generates a creatinine-blind window for the detection of GFR decline [53]. Indeed, our model predicts that with an impaired myogenic response GFR exhibits a slightly larger decrease with all treatments involving furosemide, and a significant decrease when given high drug sensitivity (Fig. 4B). The excessive decrease in GFR may lead to a corresponding decrease in urine flow. Indeed, when myogenic response is impaired, urine volume drops to the lowest level seen in all simulations following the triple treatment with high drug sensitivity (Fig. 4C).

Hydration status is also a key factor in heightening the risk of AKI (Fig. 4). ACEI and furosemide both work to reduce MAP by lowering blood volume. When water intake is not able to increase in response, blood volume, pressure, and renal blood flow can drop to dangerous levels, causing a reduction in GFR. Other factors not considered in the present study includes alterations in the glomerular ultrafiltration coefficient (e.g., in mesangial contraction [54]), reduction in oncotic filtration pressure (e.g., in proteinuria [55]), TGF resetting [56], and the presence of certain other drugs.

### 4.3 Model limitations

A significant contributing factor to triple whammy AKI that isn’t considered in the present study is nephron loss, often a consequence of hypertension. The kidney adapts to a reduction in nephron population and can, up to some critical nephron loss, maintain GFR at the normal level [57, 58]. Nonetheless, subcritical nephron loss likely elevates the risk for triple whammy AKI. To simulate kidney function under differing physiological and pathophysiological conditions, a more detailed representation of the kidney may be incorporated (e.g., for nephron function [59–61] and for renal autoregulation [62–65]).

This model is also limited in its simulation of furosemide, which focuses on the change in Na^+^ reabsorption and the resulting change in water reabsorption. However, lower water reabsorption in the loop of Henle would lead to increased pressure in the tubule. Taken in isolation, this would lower GFR and thus urine flow. These mechanics are outside the scope of this model, as the model does not account for how pressure within the nephron effects GFR but is worth considering.

In this study, we have focused on furosemide, a loop diuretic; however, other types of diuretics, such as thiazide or K^+^-sparing diuretics, are often used in treating hypertension. The main effect of any diuretic is lowering extracellular volume, which triggers the responses to sustain volemia, blood pressure, and GFR, which are inhibited by RAS inhibitors and NSAIDs. As such, we expect our results to extend to other types of diuretics.

Finally, hypertension is simulated by assuming an overactive RSNA. However, hypertension is typically a result of a variety of changes in the body, including, but not limited to, increases in systemic vascular resistance, preglomerular resistance, afferent arteriole resistance, Na^+^ retention, renin or ALD secretion, or nephron loss. Results of the present simulations will probably extend to other causes of hypertension, inasmuch as RSNA stimulates many of the other causes of hypertension, such as afferent arteriole resistance, renin secretion, and ALD secretion. Indeed, ACEI/ARB and furosemide are often prescribed to patients with heart failure and some kidney dysfunction. The cumulative effects of these drugs, as previously discussed, on feedback systems would not change, causing our results to extend to these patients, indeed, kidney dysfunction indicates an already low GFR—these patients would be especially at risk due to dehydration.

## A Appendix

This model is an application of Leete and Layton [9], which incorporates sex differences in the RAS, RSNA, and ALD into the work done by Karaaslan et al. [11] and Hallow et al. [12]. The following equations are for reference purposes, and care is taken to maintain the notation used in Karaaslan et al. [11] and Hallow et al [12], but for the sake of brevity the entire model is not reproduced here.

### A.1 RSNA

Renal sympathetic nervous activity is modeled as a baseline value (*N*_rsna_) multiplied by the effects of MAP (*α*_map_) and right atrial pressure (*α*_rap_). Equations for *α*_map_ and *α*_rap_ are taken from Karaaslaan et al. [11].

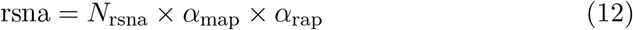

### A.2 Sodium flow past the macula densa

The filtered sodium load (Φ_filsod_) represents how much sodium is filtered through the glomerulus and equal to the plasma sodium concentration (*C*_sod_) multiplied by the GFR (Φ_GFR_).

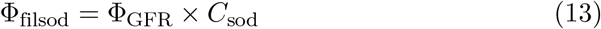

Model proximal segment tubule Na reabsorption (Φ_pt−sodreab_) is the fractional model proximal segment Na reabsorption (*η*_pt−sodreab_) multiplied by the filtered sodium load. Fractional model proximal segment Na reabsorption is the baseline value (*η*_*η*−pt_) multiplied by the effects of the filtered sodium load (*γ*_filsod_), AT1R-bound Ang II (*γ*_at_), and RSNA (*γ*_rsna_). See Karaaslan et al. [11] and Hallow et al. [12] for details of these effects.

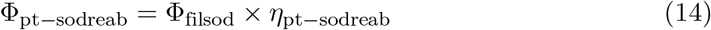

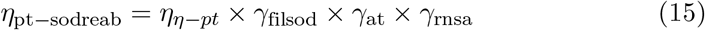

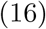

Macula densa sodium flow (Φ_md−sod_) is the filtered sodium load minus model proximal segment sodium reabsorption.

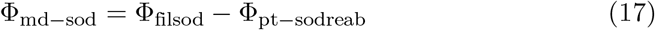

### A.3 Effects of macula densa sodium signaling

Afferent arteriole resistance is modeled as a baseline value (*R*_aa−ss_ = 32.66 mmHg/min) multiplied by the effects of rsna (*β*_rsna_), TGF (Σ_TGF_), the myogenic response (Σ_myo_), and the type 1 and 2 angiotensin II receptors (Ψ_AT1R−boundAngII−aa_, Ψ_AT2R−boundAngII−aa_ respectively).

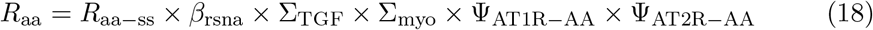

Renal blood flow (Φ_rb_) is mean arterial pressure (*P*_ma_) divided by total renal resistance (*R*_r_), the sum of afferent and efferent arteriole resistance (*R*_aa_ and *R*_ea_, respectively).

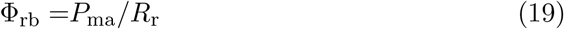

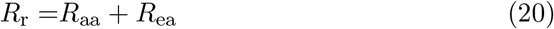

### A.4 The myogenic effect

The multiplicative effect on afferent arteriole resistance due to the myogenic effect dependent on glomerular hydrostatic pressure (*P*_gh_) and is denoted

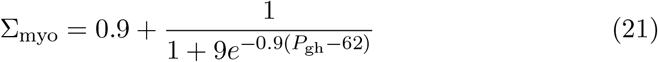

When there is high glomerular hydrostatic pressure, Σ_myo_ is high, constricting the afferent arteriole and reducing renal blood flow, thus reducing glomerular hydrostatic pressure and GFR. Similarly, when glomerular hydrostatic pressure is low, Σ_myo_ is less than 1, dilating the afferent arteriole and increasing renal blood flow, glomerular hydrostatic pressure, and GFR.

### A.5 The renin-angiotensin system

Angiotensinogen (AGT) is secreted at the rate *k*_AGT_ and is converted into Ang I through plasma renin activity (PRA). It decays with a half-life of *h*_AGT_.

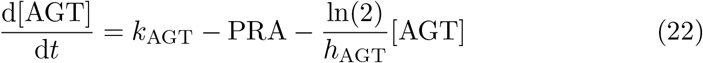

PRA is proportional to the plasma renin concentration ([PRC]) with a conversion ratio of *X*_PRC−PRA_. Renin is secreted at the rate *R*_sec_ and decays with a half life of *h*_renin_. The renin secretion rate has a baseline value of *N*_rs_ = 1 ng/ml/min and is determined by the feedback effect of AT1R-bound Ang II (*ν*_AT1_) and the effect of macula densa sodium flow (*ν*_md−sod_) described in Eq. 8.

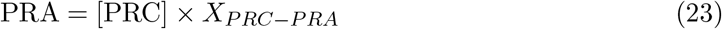

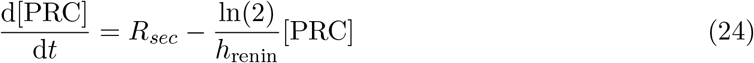

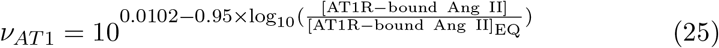

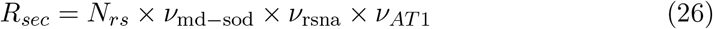

Angiotensin I ([Ang I]) is converted from AGT through PRA and is converted into angiotensin II ([Ang II]) through angiotensin converting enzyme (ACE) and chymase activity with the first order reaction rate constants *c*_ACE_ and *c*_chym_, respectively. Ang I is converted into Ang (1-7) through neutral endopeptidase activity with reaction rate constant *c*_NEP_ and decays with a half-life of *h*_AngI_.

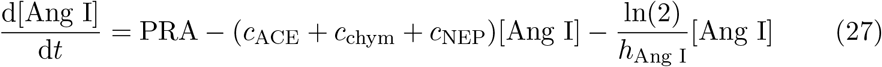

Ang II is converted from Ang I through ACE and chymase activity and is converted to Ang (1-7) through ACE2 activity and into Ang IV with reaction rate constants *c*_ACE2_ and*c*_AngII=AngIV_, respectively. Ang II binds to the type 1 and 2 receptors (AT1R and AT2R) with reaction rate constants *c*_AT1R_ and *c*_AT2R_. Ang II, AT1R-bound Ang II, AT2R-bound Ang II, Ang (1-7), and Ang IV each decay with a half-life of *h*_AngII_, *h*_AT1R_, *h*_AT2R_, *h*_Ang(1−7)_, or *h*_AngIV_, respectively.

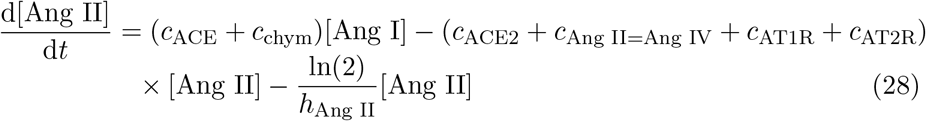

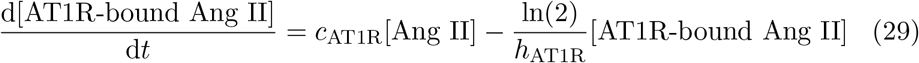

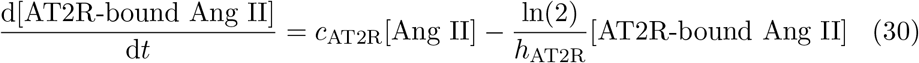

AT1R-bound Ang II and AT2R-bound Ang II each have an effect on afferent arteriole resistance (Ψ_AT1R−boundAngII−aa_, Ψ_AT2R−boundAngII−aa_ respectively).

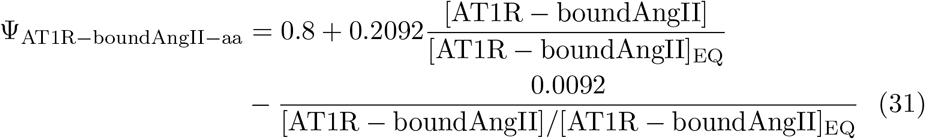

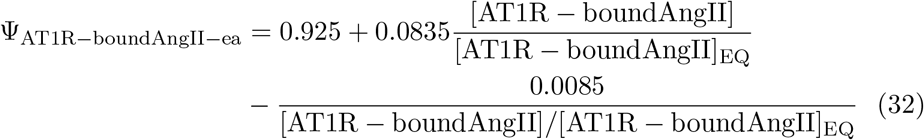

### A.6 Antidiuretic hormone

The antidiuretic hormone concentration (*C*_adh_) is modeled as the product of the normalized antidiuretic hormone concentration (*N*_adh_) times its baseline value (4 milli-units/L). Eq. 34 models the regulation mechanisms, whereby the normalized antidiuretic hormone concentration (*N*_adh_) is regulated to the normalized antidiuretic hormone secretion rate (*N*_adhs_) as done in [11]. The time constant *T*_adh_ is taken as 6 min, the baseline value of *N*_adh_ is taken as 1. The normalized antidiuretic secretion rate (*N*_adhs_) is dependent on the Na^+^ concentration (*C*_sod_) when it is above 141 mEq/L, the autonomic multiplier effect (*r*_aum_) when it is greater than 1, and the effect of right atrial pressure (*δ*_ra_). Eqs. 35 and 36 are taken from [11].

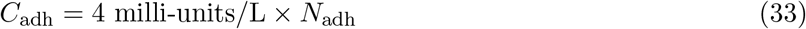

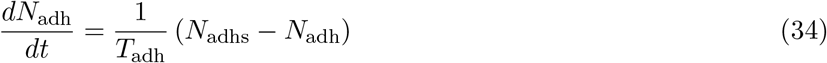

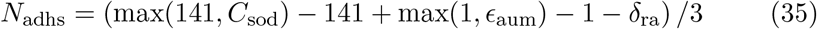

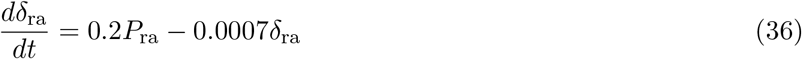

## Acknowledgments

This research was supported by the Canada 150 Research Chair program and by the Natural Sciences and Engineering Research Council of Canada, via a Discovery award (RGPIN-2019-03916).

